# Binocular Imaging with The Conformal Eyes

**DOI:** 10.1101/2022.06.06.494878

**Authors:** Jacek Turski

## Abstract

The human eye possesses a natural asymmetry of its optical components and an inhomogeneous distribution of photoreceptors causing optical aberrations and providing high acuity only at a 2-degree visual angle. Although these features greatly impact the visual system functions, they have not been supported by the geometric formulation of the fundamental binocular concepts. The author recently constructed a geometric theory of the binocular system with asymmetric eyes (AEs) integrated with the eyes’ movement to address these problems. This theory suggests that a symmetric framework can fully represent the asymmetric properties of this binocular system with the AEs. Pursuing this idea leads first to the conformal eye model furnished by the Riemann sphere for the special case of AE corresponding to the reduced eye with axial symmetry, referred to as the symmetric eye (SE) model. The conformal geometry of the Riemann sphere establishes efficient image representation by the projective Fourier transform (PFT)—the Fourier transform on the group of image projective transformations representing images covariantly to these transformations. The PFT is fast computable by an FFT algorithm in log-polar coordinates known to approximate the retina-cortical mapping of the human brain’s visual pathways. The retinotopy modeling here with PFT is compared to Schwartz’s modeling with the exponential chirp transform showing clear advantages of PFT in physiological conformity and numerical efficacy. Finally, the conformal eye model initially developed for the SE is extended to the AE. Then, in the binocular system with AEs, the PFT becomes available for image processing. The PFT combined with the conformal eye model allows binocular extensions of the previous monocular algorithms for modeling visual stability during the saccade and smooth pursuit eye movements needed to offset the eye’s acuity limitation.

## 1 Introduction

The human eye is an optical system comprising the cornea, the crystalline lens, and the nonuniform light-sensitive retina with a high acuity fovea limited to a two-degree visual angle. The human eye has an imperfect optical design: the fovea’s temporalward displacement from the posterior pole produces the eyeball’s global asymmetry, and the lens is tilted relative to the cornea. The fovea’s displacement and the asphericity of corneal surfaces are the main sources of optical aberrations. The lens’ tilt and distribution of the gradient refractive index, on the other hand, produces nearly aberration-free perception near the visual axis [38]. Moreover, the two eyes working in unison in the binocular system improve visual acuity and assist visually guided tasks with precise eye movements [16, 52].

Because our two eyes are separated laterally in the head, they receive two-dimensional disparate projections of the world. Despite this disparity, we usually perceive one world with the impression of stereoscopic depth. The basic concepts underlying binocular vision and stereopsis are retinal correspondence and horopter. For a given binocular fixation, the horopter is the locus of points in space such that each point on the horopter projects to a pair of corresponding retinal elements, one of the pair’s elements in each eye. Each pair of corresponding elements is distinguished by the quality of seeing a small object that projects onto them in a specific but subjective direction. We say that the corresponding elements have zero disparity.

Normal correspondence occurs when the fovea of one eye corresponds to the fovea of the other eye; their single visual direction is called the principal visual direction, or the Cyclopean direction. The visual directions of all other pairs of stimulated corresponding elements are perceived relative to this principal direction, the hallmark of a single binocular vision. Points far from the horopter are projected on noncorresponding elements such that they have non-zero disparity and are generally seen as double.

However, each point in the so-called Panum’s fusional region around the horopter is seen as single. Then, the brain uses the extracted disparity to create our sense of depth relative to the horopter. Further, the disparity between two spatial points in the Panum’s area provides us with stereopsis–the ability to perceive form. Even objects far from the horopter curve that we see double provide the sense of depth [24], which gives us the spatial order of objects and, hence, the phenomenal space geometry.

Recently, two binocular systems, each with a more advanced fidelity of eye model were developed. The first binocular system used the reduced eye model. In this eye model, one refractive spherical surface represents the eye optics with the fovea at the posterior pole. The nodal point and the fovea on the optical axis define the eye’s axial symmetry and are referred to as the symmetric eye (SE) model. The nodal point is 0.6 cm anterior to the eye’s rotation center. The image on the retina is impinged by the pencil of light rays based at the nodal point. The light ray directed at the nodal point passes the eye’s optical system without changing direction.

The SE model was used in the binocular system in [44] to correct the 200-year-old Vieth-Müller circle (VMC) model of the geometric horopter that is still used today in research and education. Most often, in the construction of the VMC, the nodal point is placed incorrectly in the eye’s rotation center. In this case, the analytic form of the VMC has been known for an arbitrary fixation. It follows because it remains stationary when the eyes fixate on points on the VMC. However, when the nodal point is placed at any other location, including its anatomical place, the analytic firm of the VMC was only known for symmetric fixations before 2016, see [44] where the analytic form of the geometric horopter was given for an arbitrary nodal point location and any horizontal bifoveal fixation for the first time.

The penalty for the simplicity of the VMC is that relative disparity is fixation-independent. However, the relative disparity depends on the eyes’ posture when the nodal point is in anatomical location [44]. With the eyes constantly moving even during fixations [20], this dependence should support perceptual benefits such as breaking camouflage and may also provide the aesthetic benefit of stereopsis vividly experienced by formerly stereoblind people [26].

The second binocular system used the asymmetric eye (AE) model introduced and comprehensively discussed in [47] and slightly modified in [48]. The AE extends the reduced eye with the eye’s natural misalignment of human optical elements. Based on the observed misalignment of the optical components in healthy human eyes [8, 25], the AE is comprised of the fovea’s anatomical displacement from the posterior pole by angle *α* = 5.2° and the crystalline lens’ tilt relative to the cornea by angle *β*. Angle *α* is relatively stable in the human population, and angle *β* varies between −0.4° and 4.7°. In the human eye, the fovea’s displacement from the posterior pole and the cornea’s asphericity contribute to optical aberrations that the lens tilt partially compensates for [3]. The geometric theory of the binocular system with the AEs’ developed in [48] in the framework of bicentric perspective projections advanced the work by Ogle [23] and Amigo [2] on modeling empirical horopters as conic sections. In contrast with their theory, the biologically supported horopteric conics in [48] pass through the nodal points, are integrated with eye movement, and their geometric transformations during eye movements are visualized in a computer simulation.

When eyes are binocularly fixated straight ahead such that the image planes of the AEs, defined as parallel to the lens’s equatorial planes and passing through the eye’s center, are coplanar, the constructed horopter is a straight frontal line parallel to the image planes. The distance to this fixation point is called the abathic distance and is explicitly given in [48] in terms of the ocular separation and the AE’s parameters *α* and *β*. This case of linear horopter proves that the unequal distribution of retinal corresponding points for asymmetric eyes can be defined in terms of the image plane’s symmetrical distribution of projected retinal points into the plane [48]. This geometric result suggests that one can fully dispense with the spherical retina in the AE. Making this suggestion physiologically credible and geometrically rigorous here will take us well beyond concepts typically associated with the horopter.

The paper is organized as follows. I first introduce the AE model, which defines the asymmetric binocular correspondence on the retina and the related symmetric binocular correspondence on the AE’s image plane. However, to simplify the geometric discussion, I first consider the SE model, that is, the AE with *α* = *β* = 0, emphasizing the geometric properties of the projection of the retina onto the image plane for three different locations of the nodal point: the pupil identified with the north pole of the unit sphere, the anatomical location of 0.6 cm anterior the eye’s rotation center and the rotation center.

The nodal point in the SE is often located at the rotation center, especially in computational studies; see [31] for example. However, in many publications, the nodal point has been depicted in any of the three locations mentioned above.

The north pole location of the nodal point is distinguished for providing anatomically acceptable numerical approximation and efficient tools for image processing. In particular, the projection from the retina to the image plane is the conformal stereographic mapping: it preserves the angle of the two intersecting curves under the projection. It also maps circles in the retina that do not contain the north pole to circles in the image plane. Thus, stereographic mapping preserves receptive fields’ circular shape and the retinal illuminance under the projection onto the image plane. Thus, the image plane can faithfully represent the retina.

Because stereographic mapping is not defined at the nodal point identified with the north pole, it is extended one-to-one and onto by appending to the image plane the point at infinity, which is the image of the nodal point under this mapping. The image plane with the appended point at infinity is the celebrated object in geometry and mathematical analysis and is known as the Riemann sphere [22]. Thus, the Riemann sphere gives the newest and most compelling eye model that is referred to as the conformal eye (CE).

The Riemann sphere’s conformal geometry identifies the structures of the one-dimensional complex projective geometry [6] and the Möbius geometry [14]. This identification means the group of image projective transformations resulting from the change of perspective between the scene and the eye’s orientation agrees with the group of Möbius transformations. The CE provides a unique geometric environment relevant to the intermediate-level vision computational aspects of natural scene understanding [43, 45].

Further, the Riemann sphere’s conformal geometry establishes image representation in terms of the projective Fourier transform constructed in the noncompact picture of the Fourier analysis on the semisimple group **SL**(2, C) of image projective transformations [40, 41]. This image representation changes covariantly with the image transformations induced by eye rotations. It is fast computable by FFT in log-polar coordinates known to approximate the retina-cortical mapping of the human brain’s visual pathways [42, 45]. Although the CE model is introduced and discussed for the SE model, in the last section, it is extended to the AEs.

## 2 The AE and Binocular Correspondence

The AE model is introduced and comprehensively discussed in [47] and slightly modified in [48]. The AE is shown here in a 3D drawing with some additional anatomical detail. This AE is given in Figure 1.

**Figure 1:**
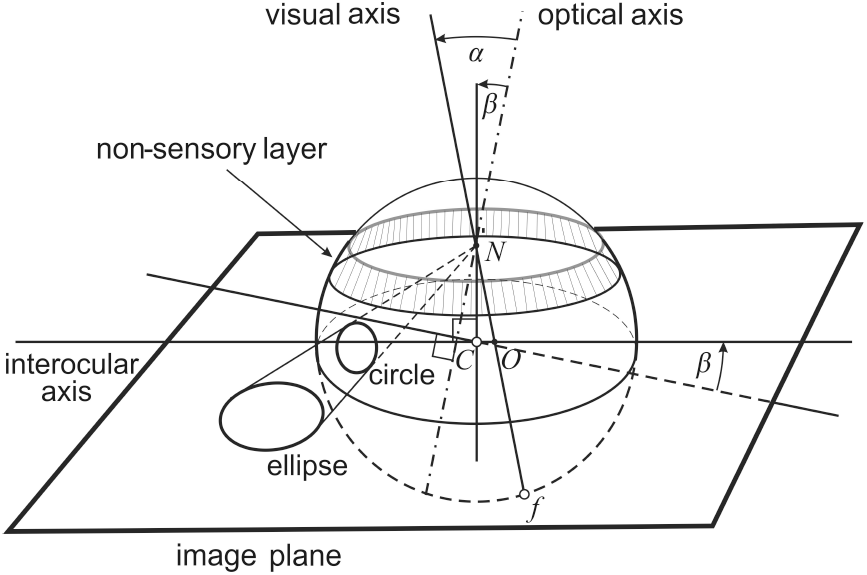
The right AE shows the non-sensory layer, whose supporting functions include the zonule—the fiber band attached to the lens that changes its curvature during accommodation [28]. The fovea *f* is displaced by angle *α* from the optical axis, and the lens is tilted by angle *β* at the nodal point *N* with both rotations in the plane spanned by the optical and interocular axes. The line through *C* and *N* is perpendicular to the lens’ equatorial plane, and the image plane is parallel to this equatorial plane and passes through the eye’s rotation center *C*, see also Figure 2. An ellipse is the image of a circle on the retina below the non-sensory layer projected into the image plane. The nodal point *N* is located at a distance |*CN* | = 0.6 cm.

The geometric theory of the binocular system with AEs fully specifies the retinal correspondence asymmetry in terms of AE’s parameters and the point of fixation. It was demonstrated in simulations of horopteric conics [48] starting from the eyes posture shown in Figure 2.

**Figure 2:**
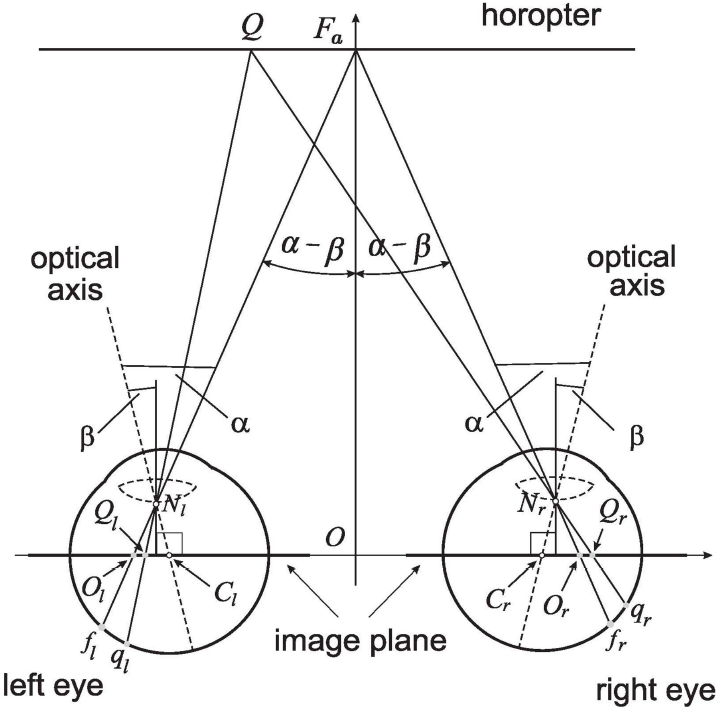
The linear horopter passing through the fixation *F*_*a*_ at the abathic distance *d*_*a*_ = |*OFa*|. In this eyes’ posture, the image planes, which are parallel to the lenses’ equatorial planes, are coplanar. The point *F*_*a*_ projects through the nodal points *N*_*r*_ and *N*_*l*_ to the foveae *f*_*r*_ and *f*_*l*_ while the point *Q* on the horopter projects to the retinal corresponding points *q*_*r*_ and *q*_*l*_ and to points *Q*_*r*_ and *Q*_*l*_ in the image plane. Although |*q*_*r*_*f*_*r*_| ≠ |*q*_*l*_*f*_*l*_|, |*Q*_*r*_*O*_*r*_| = |*Q*_*l*_*O*_*l*_| where *O*_*r*_ and *O*_*l*_ are projected foveae *f*_*r*_ and *f*_*l*_ and the equality on the image plane holds for all horizontal fixations [48].

The geometric theory in [48] shows that the non-symmetrical distribution of the corresponding retinal elements, determined by the asymmetry parameters of the AE, can be fully formulated in terms of the symmetrical distribution of their projection points in the image plane. This relationship is established by the image planes being coplanar for fixations at the abathic distance distinguished by the property that the horopter through the fixation point is a straight line parallel to the image planes, see Figure 2 and its caption for the explanation. This analysis suggests that the nodal point and the image plane encode symmetric binocular correspondence that can fully specify the AE model.

However, to dispense with the retina in the AE model, I need to show that the receptive fields and retinal illuminance are preserved under the projection from the retina to the image plane. This simple requirement will take us in the remaining part of the paper on a much longer journey through sophisticated mathematics that has striking relevance to retino-cortical image processing. Moreover, it can provide algorithms for visual stability in anthropomorphic vision systems of mobile robots.

## 3 The Conformal Geometry in Imaging

### 3.1 The Nodal Point Location and Geometry of the Image Plane

I start discussing geometries of the image plane for the AE with the condition *α* = *β* = 0 for the SE, where the optical axis and the visual axis, shown in Figure 1 for the AE, coincide. In this case, the projections through *N* of the circles on the retina representing receptive fields below the non-sensory layer are ellipses in the image plane. The ratios of the axes of the ellipses in the image plane region subtended by a visual angle of 110° at the nodal point vary from 0.7 at the high eccentricity to 1 at the foveal region. Details of the calculations are presented in Appendix A.

To discuss geometries of the image plane rigorously but also following the tradition established in biological vision by placing the nodal point at various positions. The Vieth in 1818 placed the nodal point at the eye’s pupil, Müller in 1826 placed it at the lens center, but most often, the nodal point is located at the eye’s rotation center, as originally proposed by Volkmann in 1836; see [35]. All these different locations are scattered through the contemporary research publications. However, it should be pointed out that the analytic form of the VMC as the geometric horopter has been only known for all fixations in the horizontal pane in the last case when the nodal point coincided with the eye’s rotation center. Otherwise, for all the other nodal point locations, the analytic form was known only for the symmetric fixations before the publication [44] in 2016.

I consider two different projections of the unit sphere of proximal retinal stimuli onto the image plane. The first projection, shown in Figure 3 (A), is the stereographic projection from the north pole *N* (the nodal point at the eye’s pupil) to the plane that passes through the sphere’s center *C* and is perpendicular to the line through *N* and *C*. Stereographic projection is ubiquitous in physics and mathematics and has been used for millennia in making maps. Even to mapmakers, it was well known that stereographic projection is conformal, i.e., it preserves angles of intersecting curves. Further, it maps circles in the sphere that do not contain *N* to circles in the image plane, while any circle that contains *N* projects onto a straight line. The plane is extended by upending the stereographic image of *N* such that the topology is extended by the complement of the compact sets in the plane. The extended plane with the stereographic image of *N* referred to as the point at infinity is the Riemann sphere. The resulting conformal geometry of the extended plane with its remarkable features is carefully discussed in the next sections.

**Figure 3:**
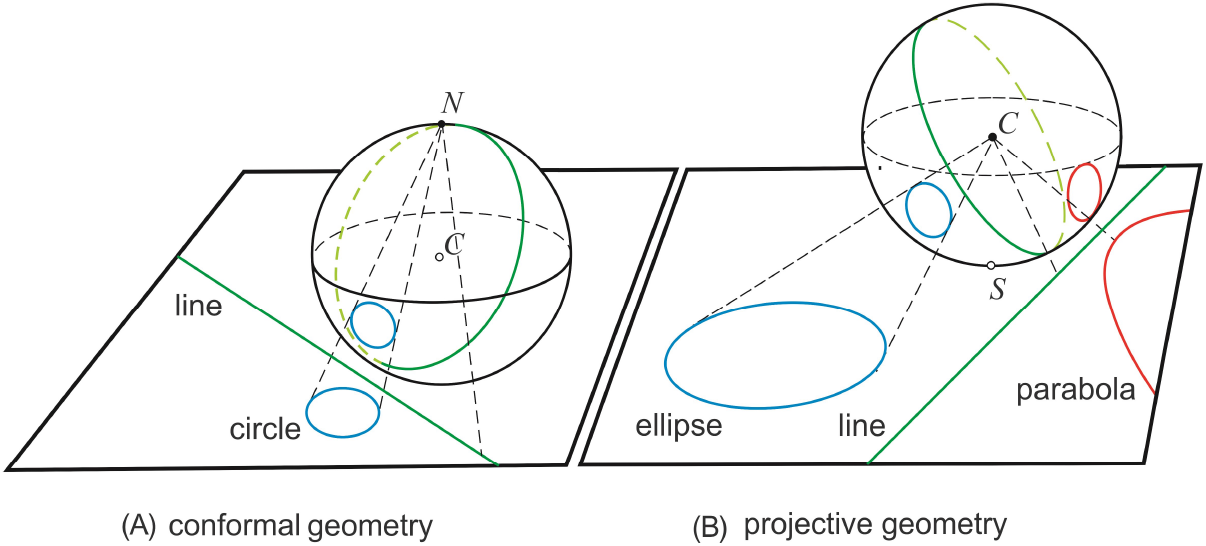
The two geometries are defined by projections of a unit sphere to the plane. (A) Conformal geometry of the unit sphere’s stereographic projection from the north pole *N* to the horizontal plane passing through the sphere’s center. A circle on the sphere that does not contain *N* projects to a circle in the plane, while a circle containing *N* projects onto a straight line. (B) Real projective geometry of the gnomonic projection from the center *C* onto the horizontal plane passing through the south pole *S*. A tangent circle to the equator is mapped to a parabola (shown), while a circle intersecting the equator in two points is mapped to a hyperbola (not shown)—the great circles other than the equator project onto straight lines.

Equally important in physics and mathematics is the sphere’s gnomonic, or central, projection from its center *C* (the nodal point at the eye’s rotation center) to the plane tangent at the south pole *S* (the fovea in the SE), shown in Figure 3 (B). The real projective plane is obtained when (1) the line at infinity is appended to the plane as the image of the sphere’s equator under the projection and (2) the sphere’s diagonal points are identified to make the projection one-to-one. Under the gnomonic projection, a circle on the sphere that does not intersect the equator is mapped to an ellipse. Otherwise, a tangent circle to the equator is mapped to a parabola (shown). In contrast, a circle intersecting the equator in two points is mapped to a hyperbola (not shown)—finally, the great circles other than the equator project onto straight lines.

Both projections discussed above are related to the two eye models that have been used in the binocular system to study geometric horopters. The stereographic projection corresponds to the nodal point identified with the eye’s pupil, which is often depicted in the literature. In contrast, the gnomonic projection corresponds to the nodal point identified with the eye’s rotation center—the commonly assumed case of the VMC in computational vision research. The results in [44], where the geometric horopters were studied for any location of the nodal point between the eye’s pupil and rotation center, show good numerical approximations of disparities near the fixation point. However, only the family of horopters for the nodal point at the pupil retains qualitative properties of the family of horopters for the anatomically correct location of the nodal point because, in both cases, the relative disparity depends on eye movements. On the other hand, the family of horopters for the nodal point at the eye’s rotation center degenerates to the family of VMCs such that relative disparity is independent of eye movements. In this case, the finest aspects of binocular vision are lost [44].

### 3.2 The Conformal Eye Model

The stereographic projection is shown in Figure 3 (A), denoted by *σ* such that *σ*(*N*) = ∞. This additional point is usually denoted by ∞ and called the point at infinity. The image plane, denoted by ℂ, is the Gaussian plane with points (*x*_1_, *x*_2_) identified with complex numbers *σz* = *x*_1_ + *ix*_2_. The extended image plane, 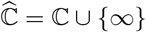, identified with the spherical retina by stereographic projection, is the celebrated object in geometry and mathematical analysis known as the Riemann sphere, [22]. The Riemann sphere is shown in Figure 4. Stereographic projection is conformal; it preserves the angle of two intersecting curves. Further, it maps circles in the spherical retina that do not contain the nodal point to circles in the image plane, while any circle that contains the nodal point projects to a straight line.

**Figure 4:**
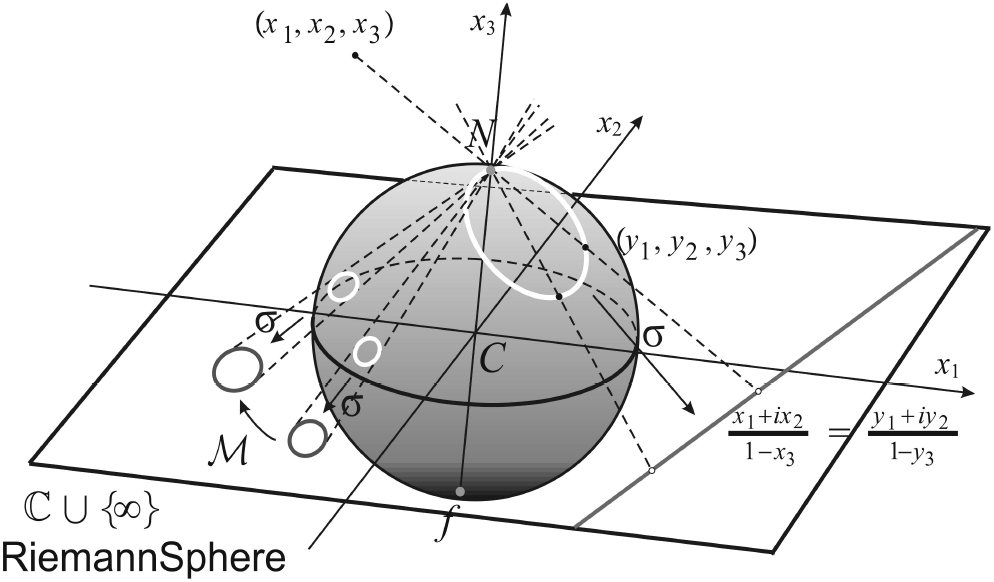
The Riemann sphere. The spatial points are projected through the sphere’s north pole *N* to the sphere and the plane. Stereographic projection *σ* maps circles in the sphere that are not passing through *N* to circles in the plane. For any two circles in the plane, there is the Möbius transformation, ℳ, that maps one circle onto the other. A circle that contains *N* is projected by *σ* to a line. Lines are regarded as circles passing through ∞.

We note that the sphere’s conformal projection to the plane, already known to Hipparchus in about 130 BC, was first named “stereographic” in 1613 by Aguilonius in *Six Books of Optics Useful for Philosophers and Mathematicians*, the treaties in which he gave the name “horopter” as well.

When the eye rotates about the eyeball center to change its gaze, simultaneously, the gaze rotates at the nodal point by the same amount, and the nodal point translates. The resulting transformation on the image plane can be decomposed into two components: (1) the translation of the image out of the image plane and stereographically projected back to the image plane and (2) stereographic projection of the image on the sphere, rotating it with the sphere, and stereographically projecting back to the image plane [39, 40, 41]. Then, as it is demonstrated in these references, the finite iterations of the transformations (1) and (2) induce the group **SL**(2, ℂ) of 2 × 2 complex matrices of determinant 1 action on the image plane by linear-fractional mappings,

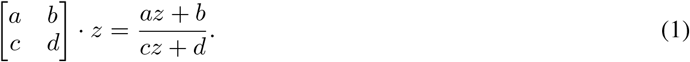

This action defines the Möbius transformations; one of these transformations, denoted by ℳ, is shown in Figure 4. However, because −*g* · *z* = +*g* · *z, g* ∈ **SL**(2, ℂ), the group of Möbius transformations is the quotient group **PSL**(2, ℂ = **SL**(2, ℂ/{±*Id*}) in which matrices ±*g* are identified [11].

Now, if *f* (*z*) is the image intensity function on the image plane, its projective transformations are the Möbius transformations that are given as follows:

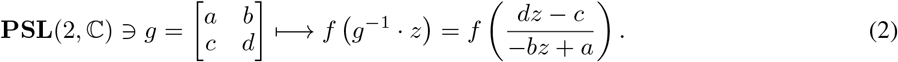

The image of a small neighborhood of any point on the sphere that does not contain *N* differs from *σ*(𝒰) in the image plane mainly by a dilation which is small for the viewing angle up to mid-periphery of about 60° and modest for the visual angle of 110° that restricts the binocular field limited by the nasal obstruction. Therefore, this conformal geometry preserves to a great degree the receptive fields and, hence, retinal illuminance as well as pixels, providing constructive properties for human vision [43, 45].

### 3.3 Geometry of the Conformal Eye

The group **SL**(2, ℂ) with the action (1) is known as the Möbius group of holomorphic automorphisms of the complex structure on the Riemann sphere 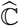 [17]. The invariants under these automorphisms furnish *Möbius geometry* [14]. Further, **SL**(2, ℂ) with the action (1) is the projective group of a one-dimensional complex geometry [6]. Thus, we have the isomorphism of the complex projective line and the Riemann sphere. This isomorphism means that the Riemann sphere synthesizes geometric and analytic (numerical) complex structures to provide a unique computational environment that is geometrically precise and numerically efficient.

The image plane does not admit a distance invariant under image projective transformations. Therefore, the conformal eye’s geometry does not possess a Riemann metric; for instance, there is no curvature. However, these transformations map circles to circles and, therefore, circles can play the role of geodesics. This fact makes the conformal model eye relevant to the intermediate-level vision computational aspects of natural scene understanding [43, 45].

## 4 The Projective Fourier Analysis

In the conformal eye model given by the Riemann sphere, the image can be represented in terms of the projective Fourier transform. The mathematical framework of this image representation is provided by Fourier analysis on the simplest semisimple group **SL**(2, ℂ), the direction in well-understood representation theory of general semisimple groups initiated by Gelfand’s school and completed by Harish-Chandra [15]—one of the greatest achievements of 20th-century mathematics.

The main idea of the Fourier analysis on groups is to decompose a space of functions defined on a set on which the group acts naturally in terms of the simplest homomorphisms of the group into the set of unitary linear operators on a Hilbert space. These simplest homomorphisms are the irreducible unitary representations of the group. In this framework, the generalized Fourier transform plays the same role on any group as the classical Fourier transform on the additive group of real numbers, where the irreducible unitary representations are homomorphisms between the additive group and the multiplicative group of complex numbers of modulus one (the circle group), given by the complex exponential functions one finds in the definition of the standard Fourier integral [30].

### 4.1 Projective Fourier Transform

The projective Fourier transform (PFT) was obtained in [40, 41] by restricting Fourier analysis on the group **SL**(2, ℂ) to the image plane of the conformal model eye. The PFT of an image intensity function, *f* (*z*), has the following expression:

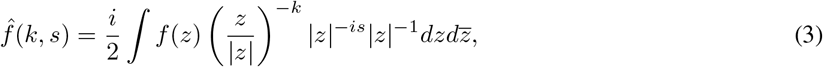

where (*k, s*) ∈ **Z** × **R**, and, if *z* = *x*_1_ + *ix*_2_, then 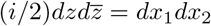. The functions Π_*k,s*_(*z*) = (*z/* |*z*|)^*k*^ |*z*| ^*is*^ in (3) play the role of complex exponentials in the classical Fourier transform; their graphs for some values of *k* and *s* are shown in Figure 5.

**Figure 5:**
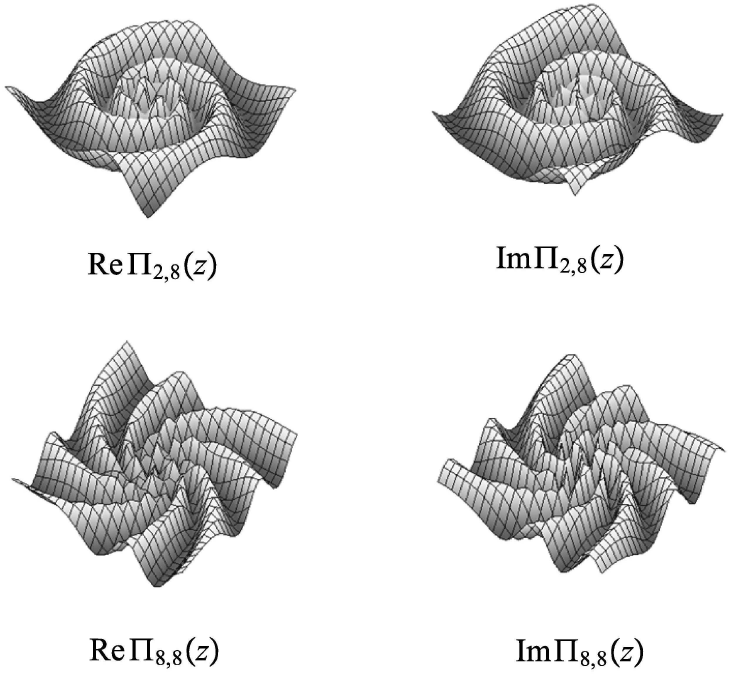
Graphs of the real and imaginary parts of the functions Π_*k,s*_ for the selected values of *k* and *s*.

In log-polar coordinates (*u, θ*) given by the complex logarithm,

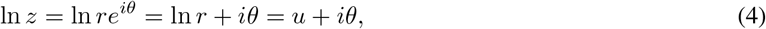

(3) takes on the form of the standard Fourier integral

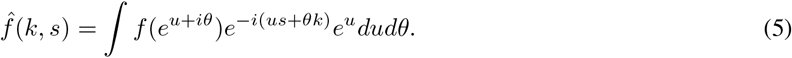

In spite of the logarithmic singularity of log-polar coordinates, an image *f* that is integrable on ℂ\{0} has finite integral (5),

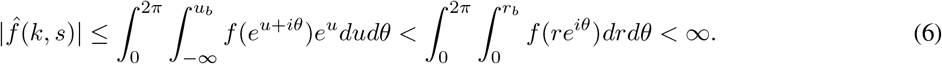

Therefore, we can remove a disk |*z*|≤ *r*_*a*_ to regularize *f* such that (*u, θ*) ∈ (ln *r*_*a*_, ln *r*_*b*_) × [0, 2*π*). This is important in the derivation of the discrete PFT.

### 4.2 Discrete Projective Fourier Transform

The PFT in log-polar coordinates, the Fourier integral in (5), can be approximated by a double (*M, N*)-Riemann sum,

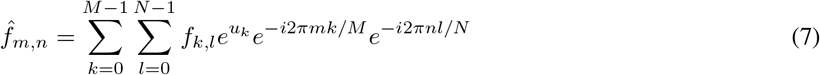

and its inverse,

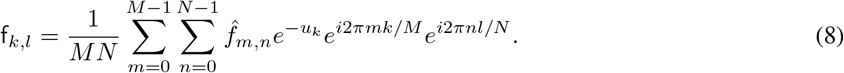

In these expressions, 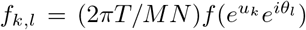 are image plane samples and f_*k,l*_ = (2*πT*/*MN*)f(*u*_*k*_, *θ*_*l*_) are log-polar samples where *T* = ln(*r*_*b*_/*r*_*a*_). I stress that 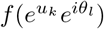 differs from f(*u*_*k*_, *θ*_*l*_); the first is defined on the image plane of the SE and the second is defined in cortical, log-polar coordinates. Both (7) and (8) can be computed efficiently by FFT.

Finally, by introducing 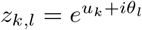 into (9) and (8) we arrive at (*M, N*)-point discrete projective Fourier transform (DPFT) and its inverse:

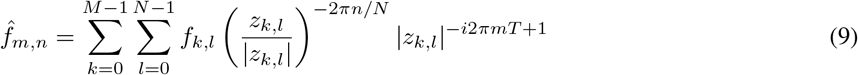

and its inverse,

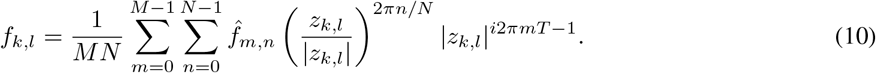

The covariant properties of the PFT are expressed as follows:

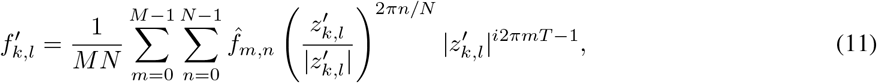

where 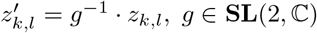 and 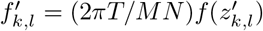.

For the covariance to image transformations demonstration on simple images, see [39, 40, 41]. However, the uniform sampling used in the discrete Fourier transform (7) is now destroyed. Therefore, for the processing of a gray level image with an FFT algorithm applied to its discrete Fourier transform (11), the nonuniform sampling theory [27] for FFT has to be used to display the transformed images. For more detailed discussion, see [41].

### 4.3 PFT and Retinotopy

How are the conformal eye model and the image representation in terms of PFT related to human vision? The human brain functions in physical space and receives information carried by light beams centrally projected onto the eyes’ retinae and transduced by photoreceptors into electrochemical signals. After initial processing by the retinal circuitry, this visual information is mainly sent to the primary visual cortex (V1), where it produces specific retino-cortical mappings and forms input to other cortical areas [50]. This immensely complex processing decodes the environment from retinal stimulation and creates a neural representation of space [33], our subjective visual space.

The log-polar coordinates (4) approximate the retino-cortical mapping of the brain’s visual pathway that topographically organizes visual and oculomotor brain areas [12]. Thus, whenever the retinal image changes as the effect of gaze rotations, the retinotopic map in V1 undergoes the corresponding changes that form the input for all next topographically organized areas that send feedback projections modulating the topographic maps.

The DPFT provides the data model for image representation that FFT can efficiently compute only in log-polar coordinates given by the complex logarithm *w* = ln *z*. As mentioned before, the central foveal region |*z*|≤ *r*_*a*_ must be removed to regularize the projective Fourier integral. The simplest, straightforward extension of the mapping *w* = *u* + *iθ*, where *u* = ln *r* with *r* > *r*_*a*_, to the region 0 ≤|*z*|≤ *r*_*a*_ can be obtained by taking linear extension of the radial part as follows

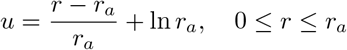

which gives the model of retinotopy required when working with the DPFT. The resulting retinotopic mapping is continuous with the continuous first derivative.

This model differs from the often used in literature retinotopy model of V1. Moreover, the superior colliculus area [32, 37] described by the mappings *w* = ln(*z* ±*a*) − ln *a* where *a* > 0 removes logarithmic singularity and ±*a* indicates, for different signs, the left or right brain hemisphere. However, both complex logarithmic mappings modeling retinotopy give similar approximations for the peripheral region. In fact, for |*z*| ≪ *a*, ln(*z* ± *a*) − ln *a* is approximately linear, while for |*z*| ≫ *a*, it is dominated by ln *z*.

In his model of retinotopy [32], Schwartz and his coworker proposed in [7] image processing in terms of the exponential chirp transform (ECT). The ECT is constructed by making the substitution:

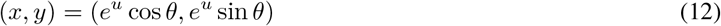

in the standard 2D Fourier integral. Because the Jacobian of the transformation is translation-invariant, this substitution makes the ECT well adapted to translations in Cartesian coordinates. There is a clear dissonance between the nonuniform retinal image polar sampling grid and this Euclidean-shift invariance of the ECT.

On the other hand, the PFT is a genuine Fourier transform constructed from irreducible unitary representations of the group of image projective transformations. Further, the change of variables by *u* = ln *r* transforms the PFT into the standard Fourier integral, which is well regularized at the logarithmic singularity. Thus, the discrete PFT is computable by FFT in log-polar coordinates that approximate the retinotopy. The difference between (12) and (4) implies that the PFT does not experience the problem of exponentially-growing frequencies, like the ECT does. Moreover, it has a band-limited original image, there is no difficulty with the Nyquist sampling condition in log-polar space, [40, 41].

Further, by the distinctive features of the complex logarithm ln *z*, ln(*e*^*iϕ*^*z*) = ln *z* + *iϕ*, ln(*ρz*) = ln *z* + ln *ρ*, the rotation and dilation transformations of an intensity function *f* (*e*^*u*^*e*^*iθ*^) expressed in exp-polar coordinates correspond to simple translations of the log-polar image f(*u, θ*) via *f* (*e*^*iϕ*^*e*^*u*^*e*^*iθ*^) = *f* (*e*^*u*^*e*^*i*(*θ*+*ϕ*^)) = f(*u, θ* + *ϕ*) and *f* (*ρe*^*u*^*e*^*iθ*^) = *f* (*e*^*u*+*v*^*e*^*iθ*^) = f(*u* + *v, θ*), where *f* and f were introduced in the previous section. These translation are very useful in the development of image identification and recognition algorithms. The Schwartz model of retinotopy, therefore, results in the destruction of these properties so critical to computational vision.

In addition, there are also psychophysiological facts that support our modeling with DPFT. The accumulated evidence points to the fact that the fovea and periphery have different functional roles in vision and very likely involve different principles underlying image processing [34, 55]. Strikingly, the authors in [34] argue that the perceptual puzzle of the curveball, when batters often report that the flight of the ball undergoes a dramatic and nearly discontinuous shift in position as the ball nears the home plate, can be explained by the difference between the foveal and peripheral processing of the ball’s image passing the boundary on the retina between these two regions.

Finally, sampling an image in log-polar coordinates conforms to the biological convergence of the retinal image sampled by photoreceptors on ganglion cells that send visual information to the visual cortex. The light carrying visual information about the external world impinged upon the retina is initially sampled by about 125 million photoreceptors. Then, after the retinal circuitry processes it, this visual information converges on about 1.5 million of ganglion cell axons that carry out the output from the eye to the brain’s visual areas. In the numerical example [42], the original image has 262, 144 pixels, whereas this image sampled in log-polar coordinates contains only 2, 664 pixels. Thus, there are about 100 times fewer pixels in the sampled image in log-polar coordinates than in the original image.

Figure 6 shows a Matlab simulation of retinotopy with DPFT. The code to perform the log-polar image transformation can be downloaded at https://www.mathworks.com/matlabcentral/fileexchange/27023-log-polar-image-sampling. Then, the standard routines of FFT and its inverse Matlab were performed; the details can be found on the Matlab webpage at https://www.mathworks.com/help/matlab/ref/fft2.html.

**Figure 6:**
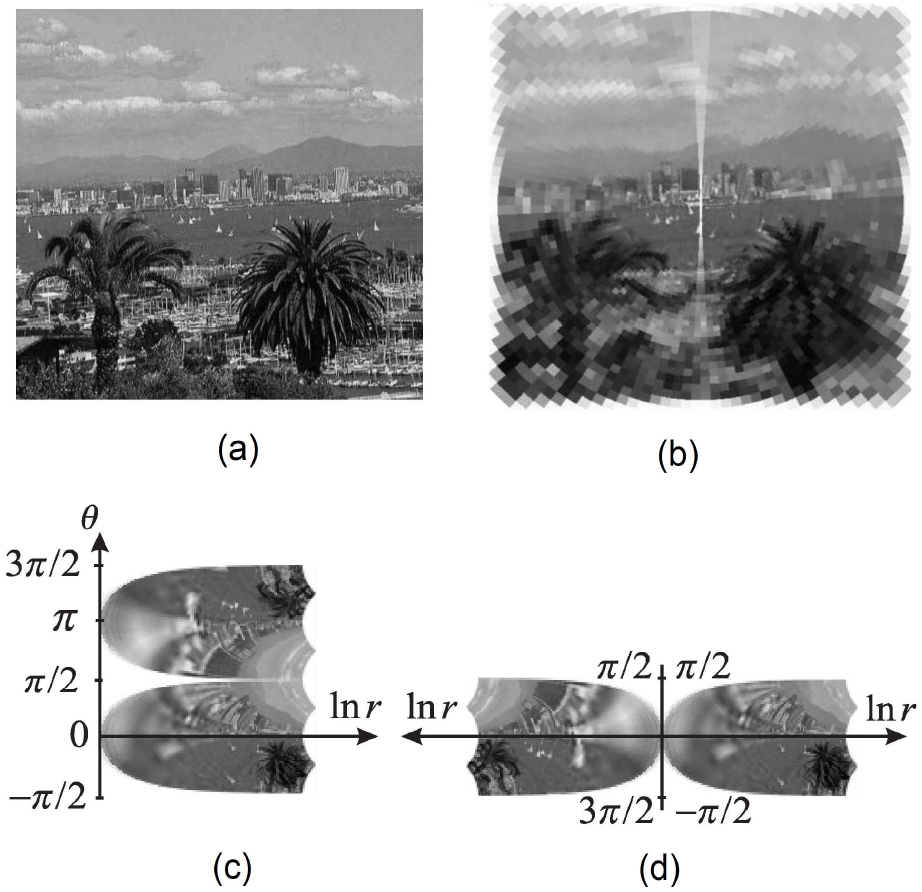
The model of retinotopy. See the text for the discussion.

In this simulation, the original San Diego harbor picture in Figure 6 (a) of the size 1024 × 1024 is resampled with 74 rings distributed exponentially and 128 sectors distributed uniformly (the exp-polar sampling). The inverse DPFT renders Figure 6 (c); both transforms, DPFT and its inverse, are computed in Matlab with FFT. Figure 6 (b) shows remapped pixel-by-pixel log-polar image in Figure 6 (c) to the Cartesian coordinates to represent the nonuniform sampling by the retina photoreceptors. The foveal region and the vertical strip are excluded from regularizing PFT, which conforms to the different image processing in the foveal and extrafoveal regions and the split theory of hemispherical image representation.

According to the split theory, the foveal region has a functional split along the vertical meridian, with each half processed in a different brain hemisphere [19]. The global retinotopy, shown in Figure 6 (d), was simulated by the cut-and-paste transformation applied to Figure 6 (c). As it was demonstrated in [42], the global retinotopy can be done with FFT in log-polar coordinates.

I should note that in a robotic camera with anthropomorphic sensors, the retinal visual information represented by Figure 6 (b) will be given by the output from the silicon retina. Then, using the DPFT representation, this output data will be processed with FFT to emphasize brightness variations in retinal illuminance and rendered, also with FFT, as the log-polar (cortical) image.

## 5 Extension to AEs

In AE, an object’s point projects to the point *z*_*β*_ on the image plane rotated by the angle *β* at the eye’s rotation center *C* rather than to the point *z* in the image plane of SE, see Figure 7. I recall that *β* specifies the tilt of the effective lens such that the image plane in AE is parallel to the lens’s equatorial plane. Because the visual axis is rotated by *α* degrees relative to the optical axis, the visual center on the image plane shifts from *C* to *O* where the visual axis intersects this plane. I need to find the transformation from *z* measured relative to *C* to *z*_*β*_ measured relative to *O*. Likewise, *y*_*β*_ is measured relative to *O*.

**Figure 7:**
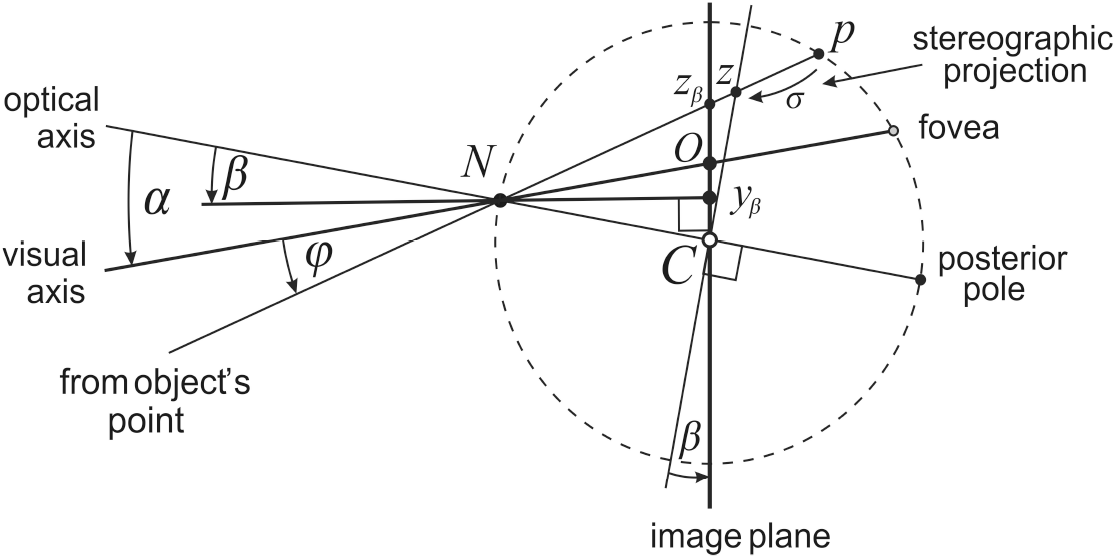
The extension of the CE model to the AE. The eyes asymmetry of optical components in the conformal eye model is represented as follows. The fovea’s displacement from the posterior point by a constant angle *α* = 5.2° gives the eyeball’s global asymmetry. The lens-cornea system is given by the image plane tilted by angle *β* from the plane perpendicular to the optical axis and the north pole *N* of the unit sphere. *β* varies between people from − 0.4° to 4.7° and *N* represents the nodal point. The imaging in this asymmetric CE is discussed in the text.

With this rule in mind, I use the right triangles Δ*Ny*_*β*_*C*, Δ*Ny*_*β*_*z*_*β*_, Δ*Ny*_*β*_*O* and Δ*NCz* to get |*Ny*_*β*_| = cos *β, z*_*β*_ − *y*_*β*_ = cos *β* tan(*α* − *β* + *φ*), *y*_*β*_ = − cos *β* tan(*α* − *β*) and *z* = tan(*α* + *φ*). Then, using these relations in *z*_*β*_ = (*z*_*β*_ − *y*_*β*_) + *y*_*β*_ and standard trigonometric identities, we obtain

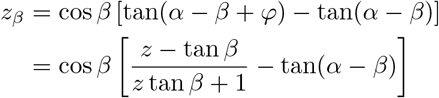

which can be written as

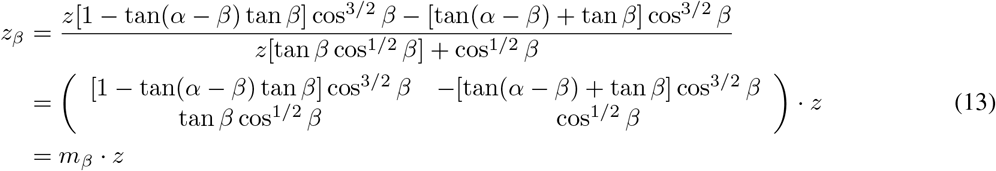

with *m*_*β*_ ∈ **SL**(2, ℂ) given in terms of AE parameters. The expressions in (13) correct the result obtained in [45] where one of terms was left out.

If *z*′ = *g* · *z*, then *z*_*β*_ = *m*_*β*_ · *z* and 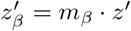 give 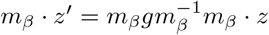 so that

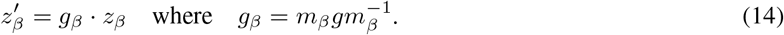

This establishes the group **SL**(2, ℂ) inner automorphism: 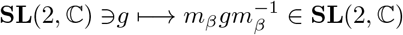 established by

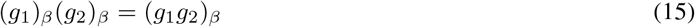

and

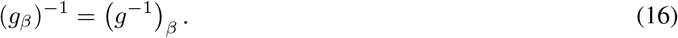

From the definition of image projective transformation, we can write,

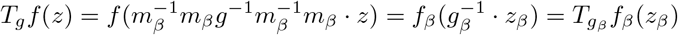

where 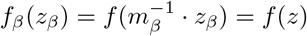. The relation between *g* and *g*_*β*_ makes the diagram

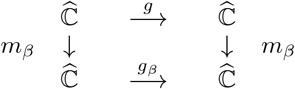

commutative such that *m*_*β*_ can be considered a coordinate transformation in the complex structure on 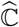. Since *g* and *g*_*β*_ have the same algebraic properties by (15) and (16), they behave geometrically in the same way. In another way of expressing the inner automorphism (14), it is to say that *g* and *g*_*β*_ are the same mappings in **SL**(2, ℂ) expressed in different coordinates (geometer’s point of view) or as the perceiving from a different perspective (physicist’s point of view).

I summarize the symmetric CE and the asymmetric CE models as follows:

1. The image projective transformations the symmetric CE

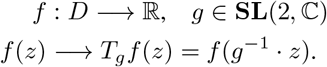
2. The image projective transformation for the asymmetric CE

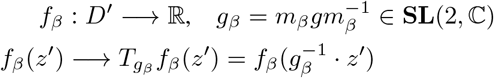

where the AE parameters *α* and *β* define coordinates change *m*_*β*_ in (13).

To conclude this section, I can say that the result of tilting and translating the image plane does not affect the asymmetric CE’s geometric and computational frameworks; this can be phrased as putting ‘conformal glasses’ on the symmetric CE.

## 6 Discussion

In [47, 48], the horopters closely resembling empirical horopters were constructed as conic sections in the binocular system with the AE model. The AE comprises two parameters: the fovea’s anatomical displacement from the eyeball’s posterior pole by angle *α* and the crystalline lens’s tilt by angle *β*. In human eyes, the foveal displacement and the cornea asphericity contribute to the eye’s optical aberrations that the lens’ tilt is partially compensating.

It was demonstrated in [48] that the non-symmetrical distribution of retinal corresponding points, due to the natural eye’s asymmetry, can be defined in the binocular system with the AEs in terms of the symmetrical distribution of the projected corresponding points into the AE’s image plane. This geometric result suggests that one can fully describe the eye imaging function with the image plane of the AE, dispensing with the spherical retina.

To this end, I first considered the AE with *α* = *β* = 0, the SE with a single refractive surface schematic eye. Following the tradition in vision science, I study three locations of the nodal point: the anatomically correct location at 0.6 cm anterior to the eye’s rotation center, the pupil’s location at the north pole of the unit sphere, and the location at the eye’s rotation center. Only for the north pole location, two crucial properties of the eye model are satisfied. First, in addition to very good approximations for anthropomorphic binocular system’s parameters, only for the nodal point placed at the north pole, it retains qualitative properties of the eye model with the correct location of the nodal point. Second, its image plane with the appended image of the north pole, i.e., the point at infinity, can be identified with the Riemann sphere. Thus, the image plane has conformal geometry. This eye model is referred to as the conformal eye.

Maybe unfamiliar to vision researchers, the automorphism group **SL**(2,ℂ) of the Riemann sphere, which provides the projective image transformations of a rotating eye, has the subgroup **SU**(2) isomorphic to the group of unit quaternions [1] that have been used for a long time in vision science [54]. Moreover, this geometry provides a unique computational environment that is relevant to the intermediate-level vision computational aspects of natural scene understanding [43, 45].

Further, the conformal eye model established image representation in terms of the projective Fourier transform [39, 40, 41]—the model eye’s intrinsic Fourier transform on the group of image projective transformations. This image representation changes covariantly with image transformations induced by eye movements. It is fast computable by FFT in log-polar coordinates that are known to approximate the retina-cortical mapping of the human brain’s visual pathways [42, 45]. The image sampling in log-polar coordinates conforms to the biological convergence of the photoreceptors sampling the retinal image on the ganglion cells that send visual information to the visual cortex with about 100 times fewer pixels than in the original image [42].

Finally, the conformal eye model established first for the SE was extended to the AE. It corrects this extension proposed in [45] for the conformal camera. Therefore, the theory in [48] that integrates binocular vision with eye movement can be supported by the Riemann sphere’s conformal geometry. Thus, this conformal AE model allows the image processing with the projective Fourier analysis developed by the author and reviewed in this article.

To conclude this discussion, I mention the future applications in the human binocular vision. The theory in [48] integrates binocular vision with eye movement. The importance of this becomes apparent when we realize that during natural viewing, the human eye’s rotational speeds during saccades are as fast as 700°/*s*, with an acceleration exceeding 20, 000°/*s*^2^ [52]. Saccadic eye movements are performed about 3-4 times/s, meaning that the brain mainly acquires visual information during 3-4 brief fixations within a second. Thus, the visual information is acquired in a sequence of discrete snapshots, but we perceive a clear and stable world. In addition, we are not only able to execute smooth pursuit eye movements that keep the foveae focused on a slowly moving object up to 100°/*s*; we also employ a combination of smooth pursuit and saccades to track an object moving unpredictably or moving faster than 30°/*s* [53, 21]. By stabilizing the tracked object’s image on the fovea, smooth pursuit eye movements (SPEMs) superimpose additional motion on the retinal images of the stationary background and the other moving objects.

The initial modeling of visual information during saccades and SPEM was carried out in [42, 46] for monocular vision. In [42], the perisaccadic predictive remapping of receptive fields by intended eye movement (via efference copy) [36] that is believed to maintain visual stability was modeled using the shift property of the projective Fourier transform in log-polar coordinates. This modeling remaps the current receptive fields to the future receptive fields before the incoming saccade takes them there. Also, it accounts for the observed in laboratory experiments the perisaccadic mislocalization of briefly flashed probes around of incoming saccade’s target [29]. In [46], in the framework of the conformal eye model, the images of the stationary background sweeping across the retina during the pursuit of a small object moving in the front of the background were obtained in the initial image plane of a discrete sequence of small rotation angle approximation of the horizontal SPAM. Thus, modeling the visual information during smooth pursuit with the conformal eye can support anticipatory image processing. This anticipation can support the stability of visual information during tracking movements by an anthropomorphic camera needed for an autonomous robot’s efficient interaction with the real world in real-time.

Thus, the basic feature underlying natural viewing is the occurrence of intricate dynamic disparities processed to maintain our clear vision that appears continuous and stable. In this regard, the theory in [48] provides the horopteric conics’ transformations during eye movement and, therefore, allows the monocular vision stability in [42] to be extended to the binocular framework. To achieve this, the theory in [48] and its extension to iso-depth conics in [49] need to include the vertical component and be integrated with 3D eye movements. I recall that the horopter is the zero-depth spatial curve. This extension will inevitably introduce a host of geometric difficulties discussed in [48].

## APPENDIX

### A Projecting Retina into Image Plane in SE Model

I demonstrate here that the projections of circles (the receptive fields) on the retina to the image plane of the SE model are conics as claimed before in Figure 1. The planar intersection of this SE model is abstracted in the following figure.

The projection from the north pole (0, 0, 1) (the eye’s pupil) of the sphere to the image plane passing through the sphere’s center is the stereographic projection. It maps circles in the sphere (the intersection of the sphere with a plane) that do not contain the north pole to the circles in the plane, see Figure 3 (A). I will demonstrate that the projections from the nodal point (0, 0, *b*) of the retina’s circles into the image plane are conic sections. Moreover, I estimate the eccentricity range of the ellipses (the ratio of their axes) in the image plane region subtended the visual angle at the nodal point of 110° for *b* = 0.6, see Figure 8.

**Figure 8:**
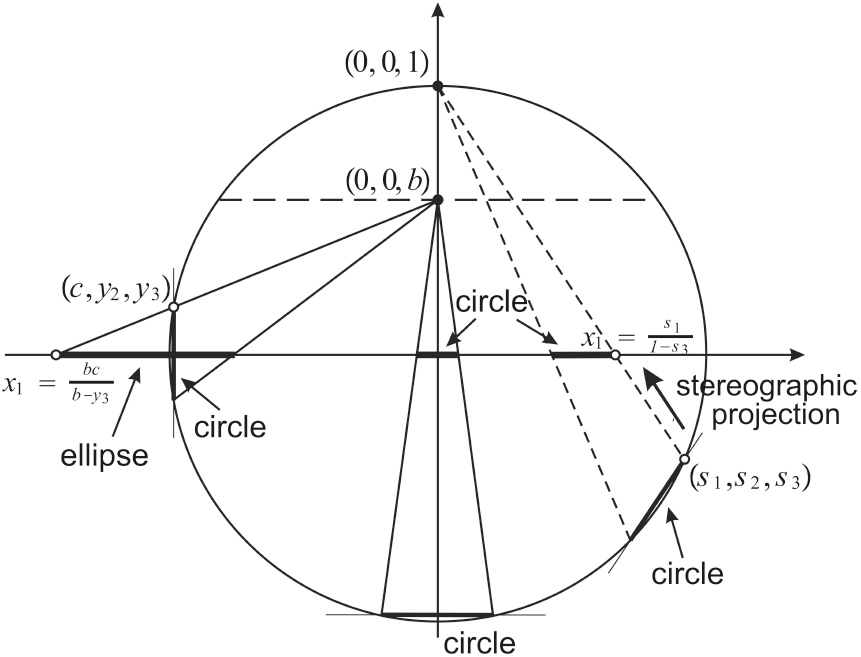
The SE model’s planar intersection. The projection through the nodal point (0, 0, *b*) from the retina to the image plane is shown with solid rays. In general, it maps circles in the retina to conic sections in the image plane. The stereographic projection is depicted with dashed rays. It maps circles in the retina (located below the horizontal gashed line) onto circles in the image plane.

The points (*c, y*_2_, *y*_3_) on the sphere 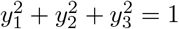 form the circle 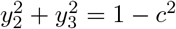. The points of the circle with *c* < 1 but close to 1 projected through (0, 0, *b*) to the image plane are

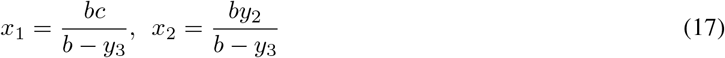

These equations are solved for *y*_2_ and *y*_3_,

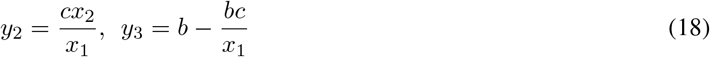

and substituted into the circle 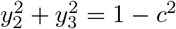 yields the equation

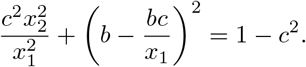

This equation can be rewritten as

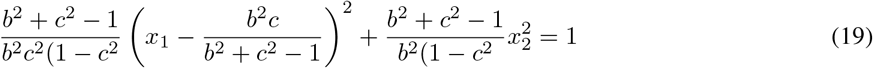

The equation (19) is an ellipse if *c*^2^ > 1 − *b*^2^ and *c*^2^ < 1, that is, if

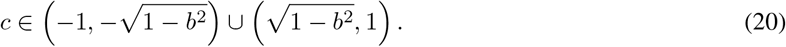

Further, the equation (19) is a parabola for 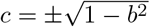 and hyperbola for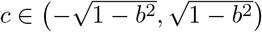.

Before I estimate the ellipses shapes, we first note that the circle 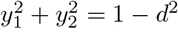 in the sphere is projected through (0, 0, *b*) to a circle in the image plane because the symmetry of projection: the center of this circle on the retina is at (0, 0, *d*) and the projection is from (0, 0, *b*), cf. Figure 8. Now, the ellipses axes ratios in the image plane region that is subtending visual angle 110° are in the interval (0.7, 1). The ration 1 is corresponding to a circle.

## Notes

### Competing Interest Statement

The authors have declared no competing interest.

### Summary of Updates

Small corrections and a new Figure 2 were added.

